# A Multi-Strain, Biofilm-Forming Cocktail of *Bacillus* spp. and *Pediococcus* spp. Alters the Microbial Composition on Polyethylene Calf Housing Surfaces

**DOI:** 10.1101/2024.11.22.624920

**Authors:** C.A. Reynolds, R.A. Scuderi, A.L. Skidmore, L. Duniere, S.Y. Morrison

## Abstract

Application of a beneficial microbial cocktail of *Bacillus* spp. and *Pediococcus* spp. was evaluated first for adherence to polyethylene calf hutch material, and second, to determine if application in situ to individual calf hutches post-cleaning influenced surface recolonization by enteric pathogens. Three treatments were utilized: 1) no application (**NC**), 2) chlorine-free, distilled water (**DW**), or 3) an application of a microbial inoculant containing *Bacillus* spp. and *Pediococcus* spp. at a concentration of 0.4 g/m^2^ of hutch space (**LF**). Thirty-six 15 × 15 cm pieces of naïve, sterile polyethylene calf hutch material received either NC or LF, were incubated at 28°C, and bacterial growth was evaluated by total aerobic plate counts at 24, 48, and 72 h post-application. Thirty polyethylene calf hutches (n = 10/treatment) were randomized to either NC, DW, or LF 24 h after cleaning. Calves were placed in the hutches 24 h after treatment application and monitored daily for 28 d. In situ surface samples were randomized by time from five unique locations within the calf hutch interior: 24 h post-cleaning, then 24 h, 7 d, 14 d, and 21 d post-application. Total aerobic plate counts and culture-independent approaches RT-qPCR and 16S amplicon sequencing were used to detect and identify the composition of the bacterial community in situ. The bacteria in the inoculant were able to successfully colonize on polyethylene, and application to individual polyethylene calf housing in situ influenced microbial diversity and reduced the presence of some undesirable bacteria on high-contact interior surfaces.

**IMPORTANCE:** Due to its multifactorial nature, neonatal calf diarrhea can be difficult to manage on farms. Clean housing environments are a critical disease control point, especially for calves less than one month of age. Application of a beneficial biofilm-forming bacterial product after cleaning of neonatal calf housing may influence the microbial communities present on the surface, particularly those that may present disease risk to calves in early life.

## INTRODUCTION

The time of highest disease susceptibility in calves is within the first few weeks of life (Urie et al., 2018). Diarrhea and other digestive issues are the leading cause of mortality in calves less than 30 d of age, with calves less than 21 d at the highest risk for infection (USDA, 2018; McGuirk, 2008; Urie et al., 2018). Morbidity caused by gastrointestinal disruption can result in high economic loss due to treatment costs, impaired growth, diminished reproductive efficiency, and poor lactation performance (Cho and Yoon, 2014; Urie et al., 2018). Early-life stressors of diarrhea or antimicrobial treatments can negatively affect the diversity and stability of the intestinal microbial population and gut barrier function, which can increase the susceptibility of calves to disease, interfere with intestinal permeability, and impede nutrient absorption (Araujo et al., 2015, Steele et al., 2016; Malmuthuge and Guan, 2017; Tao et al., 2020). The multifactorial causes of diarrhea (bacterial, viral, and protozoal pathogens) make it challenging to manage due to variations between farms in management practices, housing, feeding, and environmental factors (Barrington et al., 2002; Maunsell and Donovan, 2008). Therefore, it is important to recognize the contribution of management practices to calf health (Wells et al., 1996).

The health of pre-weaned calves can be influenced by factors such as colostrum quality and management, hygiene, bedding, and proper cleaning of pens and feeding equipment (Godden, 2008; Renaud et al., 2018; Barry et al., 2019). Causative agents of diarrhea after seven days of age are most often found in the calf housing environment (McGuirk, 2008). The cornerstone of disease control measures is proper cleaning and disinfection procedures, which can remove up to 100% of surface bacterial load if performed correctly (Barrington et al., 2002). Superficial imperfections harbor bacteria that can evade a cleaning and disinfection process, which then allows them to persist on or re-colonize surfaces (Latorre et al., 2010).

Most bacteria form biofilms as a survival and proliferation mechanism, utilizing an extracellular polysaccharide matrix as a means of adherence and protection from cleaning agents, antimicrobials, and environmental assaults (Srinivasan et al., 2021), and 90% of all bacterial cells on earth exist within a biofilm (Flemming et al., 2016; Flemming and Wuertz, 2019). Biofilms are extremely common in the dairy industry, particularly in milk processing and storage equipment and animal cooling systems (Gopal et al., 2015; Shpigel et al., 2015; Shemesh and Ostrov, 2019), though currently little is known about the nuances of microbial communities in livestock housing environments (Guéneau et al., 2021).

With the intensified need for judicious use of antimicrobials in food animal species dictating and encouraging the shift to preventative strategies rather than treatment of disease (Smith, 2015), certain antagonistic and bioactive effects of beneficial biofilms may promote a OneHealth approach to disease management (Guéneau et al., 2022b). Biofilm-forming species of non-pathogenic bacteria could be used as competitive inhibitors against undesirable organisms by dominating the surface ahead of contamination (Vlamakis et al., 2013), or by disrupting the cellular mechanisms required for pathogenicity (Tazehabadi et al., 2021). Microbial-based applications have been previously studied in high-density swine and poultry production environments. Application of a beneficial biofilm solution containing *Bacillus* and *Pediococcus* spp. at a rate of 0.2 g/m^2^ following a cleaning and disinfection procedure reduced presence of Enterobacteriaceae and Enterococcaceae families in in poultry production buildings after animals were re-introduced (Guéneau et al., 2022a) and an application of *Bacillus* spp. products has been evaluated in piglet nurseries as a comparison to cleaning and disinfection, but did not have the same competitive exclusion power against *Escherichia coli*, *Enterococcus* spp., fecal coliforms, and methicillin-resistant *Staphylococcus aureus* if not coupled with a cleaning and disinfection procedure (Luyckx et al., 2016). To our knowledge, no study has yet evaluated the use of beneficial biofilms in individual pre-weaned calf housing. However, the success of this product in swine and poultry facilities suggests its applicability to the pre-weaned calf environment.

The two objectives of this study were to assess beneficial bacterial growth and adherence to polyethylene calf hutch material, and second, to evaluate if the application of a beneficial biofilm solution to individual calf housing after cleaning influenced surface re-colonization by enteric pathogens. It was hypothesized that the polyethylene material used to construct calf hutches would be a suitable surface for bacterial growth and that a reduction of enteric pathogens on the calf hutch surface would be observed in hutches treated with the beneficial biofilm solution compared to untreated.

## MATERIALS AND METHODS

The study was conducted from 2021-06-14 to 2021-07-16 at the William H. Miner Agricultural Research Institute (Chazy, NY). All experimental procedures involving calves were approved by the Animal Care and Use Committee of the William H. Miner Agricultural Research Institute (2021AUR03).

### Preparation and Handling of Treatments

Treatments included 1) a negative control (**NC**; no application) 2) a positive control application of distilled water (**DW**), or 3) an application of a microbial inoculant containing six biofilm-forming strains of *Bacillus spp.* and two strains of *Pediococcus spp.* (**LF;** LALFILM PRO, Lallemand SAS, Blagnac, France). Sachets of the inoculant were provided by the manufacturer and were stored at 20°C until use. The concentration of LF for use in the calf environment was 0.4g/m^2^ of hutch space for both laboratory and in situ experiments, achieved by rehydration with distilled water. Treatments were prepared approximately 30 min before application. At the manufacturer’s recommendation, opened sachets of LALFILM PRO were vacuum-sealed and stored at 4°C until the next use.

### Laboratory Confirmation of Biofilm Formation

Thirty-six 15 × 15 cm pieces of naïve, polyethylene calf hutch material (Calf-Tel; Hampel Corp., Germantown, WI) were autoclaved on a dry cycle (121°C, 15 min), manually cleaned using a scrub brush and non-chlorinated detergent (Prefer; IBA Dairy Supplies, Millbury, MA), and allowed to dry inside of a biosafety cabinet at 20°C for 24 h. Following the 24 h dry time, one side of each hutch piece (n = 6 pieces/treatment/time) randomly received an application of either NC or LF. A mark was made on the pieces to identify the side with the treatment application. A 1 L handheld spray bottle (Ace Hardware, Oak Brook, IL) with an adjustable spray nozzle was used to evenly apply the LF treatment to the hutch pieces without allowing droplets to form or the solution to run. The hutch pieces were suspended vertically inside an incubator and allowed to incubate for 24-72 h at 28°C.

### Sampling of Hutch Pieces

Six hutch pieces per treatment (NC and LF) were randomly identified for sampling at 24, 48, and 72 h post-application. The pieces were sampled promptly after removal from the incubator with one half of a 3M Sponge-Stick (3M; St. Paul, MN) dampened with 10 mL sterile saline cell suspension buffer, and the entirety of the 15 × 15 cm square was sampled.

### Application of Beneficial Biofilm Solution In Situ

Thirty individual, 3.3 m^2^ polyethylene calf hutches (PolyDome Products, Litchfield, MN) were cleaned using a long-handled brush, non-chlorinated detergent, and a high-pressure washer set to provide a constant rinse water temperature of approximately 60°C (HDS; Kärcher North America, Aurora, CO). After cleaning, hutches were removed from the wash area and allowed to dry for 24 h. Hutches were blocked by date of cleaning and arranged by block in parallel rows to allow for similar exposure to sun, wind, traffic, and precipitation. Hutches were placed atop crushed stone pads, bedded with fresh, kiln-dried sawdust after the 24 h drying time, and randomized within block to receive either NC, DW, or LF (n = 10 hutches/treatment). The LF and DW treatments were applied after the hutches were bedded using 2 L pressurized hand sprayers with adjustable spray nozzle settings (Chapin Manufacturing, Batavia, NY), each expelling spray at a similar rate. Treatments were applied uniformly from the top down (starting at the ceiling), beginning from the rear of the hutch, and moving to the front to ensure minimal overlap of application areas and without allowing droplets to form or the solution to run. After treatment applications, all hutches (regardless of treatment) were allowed to dry for 24 h.

All research staff were blinded to treatment assignments, and all treatment applications were performed by a single person to reduce the variability of treatment application.

### Management and Monitoring of Calves

Calves of similar age within block (17.1 ± 12.9 days of age at enrollment) were placed in the hutches 24 h post-application and fed, managed, and treated as per normal farm procedure. Bedding in the hutches was replenished as necessary throughout the treatment period. To monitor for potential adverse effects of the product, individual health scores for each calf were recorded daily between 0800-1000 h using the University of Wisconsin Calf Health Scoring Chart (McGuirk, 2015) for 28 d. Calves’ fecal, respiratory, and hydration scores were recorded daily by the same trained individuals, and scored on a 1-4 scale (1= normal; 4= severe illness).

### Environmental Conditions

Temperature and relative humidity of the calf environment were measured every 10 min daily from 2021-06-14 to 2021-07-16 with a data logger (HOBO U23 Pro v2 Temperature/Relative Humidity Data Logger, Onset Computer Corporation, Bourne, MA) placed in a protective housing located at hutch height.

### Sampling of Calf Hutches

Surface samples were collected from each hutch using one half of a 3M Sponge-Stick (3M; St. Paul, MN), dampened with 10 mL of sterile saline cell suspension buffer. One 15 × 15 cm sample was collected at random from one of five unique surfaces in the hutch interior at 24 h post-cleaning, prior to treatment application; 24 h post-treatment application prior to animal entry; then 7, 14, and 21 d post-treatment application. Each location was sampled once per hutch, per timepoint for a total of 150 samples. To minimize variability, a single person performed all sampling. Samples were placed on ice for transport to the laboratory for analysis.

### Cell Recovery, Enumeration, and DNA Extraction

Total aerobic cell counts on non-specific media were performed according to Stone et al. (2020) for samples collected from the polyethylene hutch pieces and from the hutches, with modifications to the volumes and buffers. Each sponge was aseptically removed from the stick, manually wrung out into a sterile petri dish, and rinsed with 15 mL of sterile saline buffer to recover cells. The sponge was wrung out again, and the recovered solution was centrifuged at 6,888 × *g* for 35 min at 4°C. The resulting cell pellet was then re-suspended in 1 mL of buffer, of which 100 µL was used for bacterial enumeration. Following serial dilutions, 100 µl of solution from each serial dilution was spread onto TSA plates (Becton Dickinson, Franklin Lakes, NJ) in triplicate, and cell enumeration was performed after incubation at 28 ± 2°C for 24-72 h.

For the in situ samples, the remaining 900 µL of recovered solution was re-centrifuged under the same conditions for 15 min. This cell pellet was then re-suspended in 750 µL of BashingBead Buffer (Zymo Research, Irvine, CA) in a sterile microcentrifuge tube, and stored at -80°C until DNA extraction. Extraction of microbial DNA from the in situ samples was performed using a Zymo Quick-DNA Fecal/Soil microbe kit (Zymo Research, Irvine, CA), following the manufacturer’s instructions. DNA in each of the samples was quantified using a NanoDrop 2000 spectrophotometer (Thermo-Fisher Scientific, Waltham, MA), then stored at -80°C until further analysis using real time qPCR and 16S amplicon sequencing.

### Detection of Environmental Enteric Pathogens and Lactic Acid Bacteria with RT-qPCR

Microbial DNA isolated from samples in situ was analyzed using RT-qPCR for detection of *Escherichia* spp.*, Salmonella* spp*., Cryptosporidium parvum,* and total lactic acid bacteria (LAB), and the parameters are outlined in Supplementary Table 1. Briefly, each primer pair assay for *Escherichia* spp.*, Salmonella* spp*., and Cryptosporidium parvum* was optimized using SYBR Green fluorescent dye in 25 µL reaction volumes containing 12.5 µL of SYBR SsoAdvanced SYBR Green Supermix (BioRad Laboratories, Hercules, CA), 5.5 µL of sterile milliQ water, 2.5 µL of forward and reverse primers, and 2.0 µL of template DNA using a CFX96 C1000 thermal cycler (BioRad Laboratories, Hercules, CA). Total lactic acid bacteria (LAB) were quantified using 16S consensus primers, which have been previously used in fecal samples for the detection of *Lactobacillus*, *Leuconostoc*, and *Pediococcus* (Furet et al., 2009). Amplification was similarly performed in 20 µL reaction volumes containing 10 µL of SYBR SsoAdvanced SYBR Green Supermix (BioRad), 4µL of sterile milliQ water, 2µL of forward and reverse primers, and 2µL of template DNA. Standards, non-template controls (sterile milliQ water), and positive controls were analyzed in triplicate for each assay whereas unknown samples were analyzed in duplicate. After each assay, the efficiency was evaluated using the resulting standard curve (efficiency = 10^(-1/ Slope)^-1)*100). The mean starting quantity of target DNA from each sample was used to calculate the number of gene copies per DNA extracted from swab samples (expressed as µg of DNA per cm^2^), and a threshold of 3 log_10_ copies was used as a limit of detection.

### 16S Sequencing and Bioinformatics

To measure bacterial diversity and taxonomic composition of microbial communities in situ, DNA samples were also analyzed by 16S rRNA gene amplicon sequencing. The hypervariable V3-V4 regions of the 16S rRNA gene were targeted using the primers set 341F 5’-CCTACGGGAGGCAGCAG-3’ and 806R 5’-GGACTACNVGGGTWTCTAAT-3’ (1,2). A total of 150 samples were sequenced on a MiSeq high-throughput sequencer (Illumina, San Diego, CA) at the GeT-PlaGe facility (INRAE, Genotoul, Toulouse, France). MiSeq Reagent Kit v3 was used according to the manufacturer’s instruction (Illumina Inc., San Diego, CA) and with paired-end 250 bp sequences were generated. The 16S rRNA gene sequences were processed with the DADA2 pipeline 1.22.0 (3) for the pipeline’s steps of filtering, trimming, dereplication, sample composition inference and chimera removal. The Silva nr v.138.1 database (4) was used to assign taxonomy to the Amplicon Sequence Variances (ASV) with a minimal bootstrap of 50. The sequencing run produced a total of 4,422,705 reads that were used as input for bioinformatic analysis and a total of 3,059,006 reads were kept after trimming, chimeral removal and filtration steps. Rarefaction curves were analyzed to confirm the correct depth sequencing of each sample. Each treatment was designated as a unique group for the multi-pattern analysis using 999 permutations for each group.

## STATISTICAL ANALYSIS

Thirty-six hutch pieces and thirty hutches were included in the final datasets. Data from the laboratory study was analyzed as a completely randomized design, and the in situ study was analyzed as a randomized, complete block design using the Statistical Analysis System (SAS) v. 9.4 (Cary, NC). Temperature and relative humidity of the calf hutch environment were analyzed using the MEANS procedure of SAS and are reported as descriptive statistics (mean ± standard deviation). Individual daily health scores for calves were summarized but not analyzed in detail, as the study was not sufficiently powered to detect health effects.

Data was tested for normality using the UNIVARIATE procedure and homogeneity of variance tested using Levene’s test within the GLM procedure of SAS. Data from the laboratory and in situ experiments were analyzed using PROC GLIMMIX. The model for the laboratory experiment included hutch piece as experimental unit and fixed effects of treatment, time, and their interaction. The in situ ANCOVA model for total aerobic cell counts, qPCR data, and LAB ratio analysis included hutch as the experimental unit, fixed effects of treatment, time, and their interaction, and block as a random effect.

Alpha-diversity and top 20 ASVs from the 16S sequencing results were analyzed using PROC MIXED, with hutch as the experimental unit and fixed effects of treatment, time, and their interaction and block as a random effect. For variables with repeated measures, a covariate (post-cleaning sampling timepoint) was used to analyze the data to determine changes over time, with hutch within treatment as the subject. The covariance structures compound symmetry, simple, unstructured, and autoregressive order 1 were tested, and whichever resulted in the lowest Akaike’s information criterion value was used (Littell et al., 1996).

Sequencing and bioinformatics data were uploaded into R for analysis using the Phyloseq 1.42.0 R package (5) as similarly described by Duniere et al. (2024). Data from the Phyloseq object was used for diversity analysis using rarefied data for alpha diversity and transformed data for beta-diversity analysis using the variance stabilization transformation in DESeq2. The Unifrac distances were evaluated by principal coordinates analysis (PCoA) and further analyzed by permutational multivariate analysis of variance using distance matrices (PERMANOVA) using the Adonis approach in the R package Vegan 2.6-4. Alpha diversity results and the relative abundances for the most abundant ASV were exported to Excel files for statistical analysis. Additionally, the IndicSpecies package in R (1.7.14) was used to identify indicator species at the genus level using the multi-pattern analysis test (Duniere et al., 2024).

In this experiment, the total number of ASV’s only being identified in the tested group (A value), prevalence of the ASV across samples within the tested group (B value), and the statistical significance of the relationship within that group expressed as a *P*-value were reported. The statistical strength of relationships determined through a multipattern analysis are then evaluated using a permutation test. Relative abundance figures were generated at the genus level using the filtered data for the 20 most abundant ASVs, which were expressed as a percent of the total reads per sample.

Least squares means were reported and Tukey-Kramer adjustment for multiple comparisons was analyzed when a significant treatment by time effect was observed. Significance was declared at *P* ≤ 0.05 and tendencies reported at 0.05 < *P* ≤ 0.10.

## RESULTS

Bacterial growth was confirmed on the LF hutch pieces 24 h post-application (Table 1; 4.36 ± 0.00 log_10_ CFU/cm^2^; Treatment, *P <* 0.001). As expected, bacterial growth was below the limit of detection in samples taken from the NC pieces. Cell counts of the LF-treated pieces decreased between 24 and 48 h post-application (4.36 and 3.61 ± 0.01 log_10_ CFU/cm^2^, respectively) and tended to be different between 48 and 72 h (3.61 and 3.56 ± 0.01 log_10_ CFU/cm^2^, respectively).

**Table 1.** Total aerobic cell counts (log_10_ CFU/cm^2^) over 72 h enumerated from 15 × 15 cm polyethylene calf hutch pieces treated with a negative control (NC; no application) or a beneficial microbial cocktail containing *Bacillus spp.* and *Pediococcus spp*. at a concentration of 0.4 g/m^2^ (LF).

| Treatment<sup>1</sup> | Overall mean | Time Post-Application |  |  | SE | P-values |  |  |
| --- | --- | --- | --- | --- | --- | --- | --- | --- |
|  |  | 24 h | 48 h | 72 h |  | Treatment | Time | Treatment × Time |
| NC | <LOD<sup>2</sup> | <LOD | <LOD | <LOD | 0.01 | <0.001 | <0.001 | <0.001 |
| LF | 3.85<sup>a</sup> | 4.36<sup>a</sup> | 3.61<sup>bx</sup> | 3.56<sup>by</sup> |  |  |  |  |

<sup>1</sup>Treatments were a negative control (NC; no application) or a beneficial microbial cocktail (LF; Lallemand SAS, Blagnac, France) at a concentration of 0.4 g/m<sup>2</sup>.
<sup>2</sup> Results were below the limit of detection (LOD).
<sup>ab</sup> Means within each row with differing superscripts were significantly different ( $P \leq 0.05$ ).
<sup>xy</sup> Means within each row with differing superscripts tended to be different ( $0.05 < P \leq 0.10$ ).

There was a significant Treatment × Time interaction (*P* = 0.003) for in situ total aerobic cell counts (log_10_ CFU/cm^2^) shown in Table 2. Total aerobic cell counts (log_10_ CFU/cm^2^) were not different between treatments at 24 h or 7 d post-application. However, cell counts were greater in the DW-treated hutches at 14 d post-application compared to NC (4.74 and 3.55 ± 0.25 log_10_ CFU/cm^2^, respectively) and total aerobic cell counts (log_10_ CFU/cm^2^) tended to be greater in LF-treated hutches compared to NC (4.44 and 3.55 ± log_10_ CFU/cm^2^, respectively), but no differences were noted between DW and LF at this timepoint. As expected, the aerobic bacterial load of the hutches increased over time (*P* < 0.001). It is interesting to note that a numeric increase of nearly one full log_10_ CFU/cm^2^ was observed in the DW-treated hutches through the duration of the treatment period, particularly between d 7 and d 14 (3.58 to 4.74 ± 0.27 log_10_ CFU, respectively). Increases in aerobic cell counts were also observed in the NC-treated hutches between d 14 and 21 (3.55 to 5.58 ± 0.29 log_10_ CFU/cm^2^). However, no increases in log_10_ CFU/cm^2^ were observed in the LF-treated hutches at any of the sampling timepoints.

**Table 2.** Total aerobic cell counts (log_10_ CFU/cm^2^) enumerated in situ over a 21-d treatment period from polyethylene calf hutch surfaces treated with a negative control (NC; no application), a positive control of distilled water (DW), or a beneficial microbial cocktail containing *Bacillus spp.* and *Pediococcus spp*. at a concentration of 0.4 g/m^2^ (LF).

| Treatment<sup>1</sup> | Overall mean | Time Post-Application |  |  |  | SE | P-values |  |  |
| --- | --- | --- | --- | --- | --- | --- | --- | --- | --- |
|  |  | 24 h | 7 d | 14 d | 21 d |  | Treatment | Time | Treatment × Time |
| NC | 3.93<sup>y</sup> | 3.27 | 3.32 | 3.56<sup>by</sup> | 5.58 | 0.10 | 0.08 | <0.001 | 0.003 |
| DW | 4.13 | 2.99 | 3.58 | 4.74<sup>a</sup> | 5.19 |  |  |  |  |
| LF | 4.29<sup>x</sup> | 3.72 | 3.80 | 4.43<sup>abx</sup> | 5.23 |  |  |  |  |

<sup>1</sup> Treatments were a negative control (NC; no application), a positive control of distilled water (DW), or a beneficial microbial cocktail (LF; Lallemand SAS, Blagnac, France) at a concentration of 0.4 g/m<sup>2</sup>.
<sup>ab</sup> Means within each column with differing superscripts were significantly different ( $P \leq 0.05$ ).
<sup>xy</sup> Means within each column with differing superscripts tended to be different ( $0.05 < P \leq 0.10$ ).

Total copy numbers of gene targets for *Escherichia* spp., *Salmonella* spp., and *C. parvum,* and total lactic acid bacteria amplified through RT-qPCR are expressed in Table 3. Targets for *Escherichia* spp., *Salmonella* spp., and *C. parvum* were all below the limit of detection of 3 log_10_ copies/cm^2^ irrespective of treatment, so while the results are reported, inferences on microbial interactions and effects of treatment are limited. Copy numbers of gene targets for LAB, which were above detection limit, were greater in LF-treated hutches than in NC or DW for the duration of the treatment period (Treatment × Time, *P* < 0.001). As expected, significant increases in LAB gene targets were detected in LF hutches at 24 h and 7 d post-application compared with NC and DW and tended to be reduced compared with DW at 14 d post application.

**Table 3.**
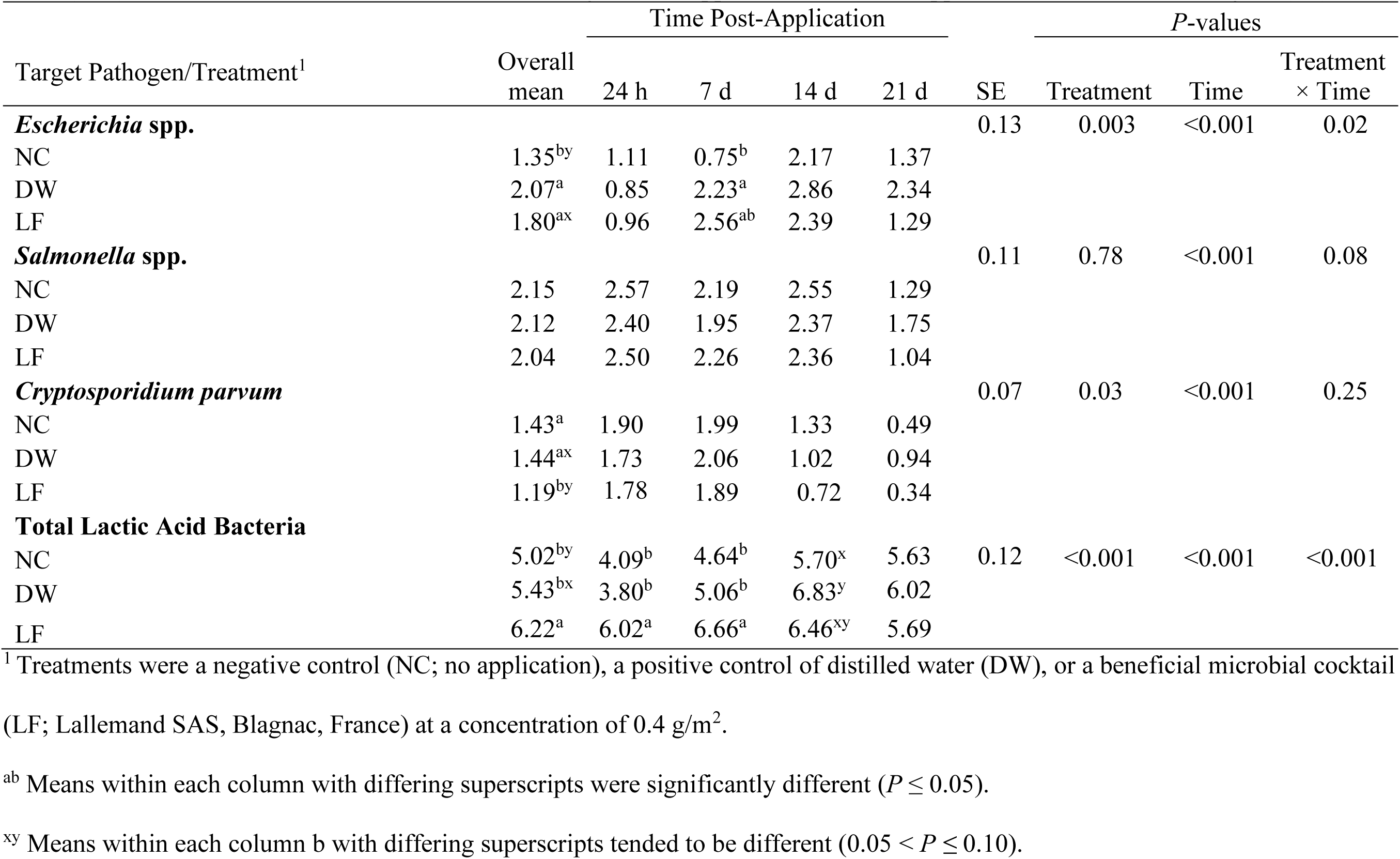
DNA concentrations (log µg DNA/cm^2^) of selected enteric pathogens and total lactic acid bacteria detected in situ over a 21-d treatment period on polyethylene calf hutch surfaces treated with a negative control (NC; no application), a positive control of distilled water (DW), or a beneficial microbial cocktail containing *Bacillus spp.* and *Pediococcus spp*. at a concentration of 0.4 g/m^2^ (LF).

Bacterial species present on hutch surfaces and alpha-diversity indices were all affected by treatment (Table 4; *P* < 0.05). The lowest number of observed species overall was observed in LF compared with DW and NC (91, 108, and 123, respectively), with significant reductions of observed species in LF treated hutches compared with NC (95 vs 161 ± 6, respectively) at 14 d post-application. At 24 h post application, Shannon diversity indices tended to be lower in LF treated hutches compared with NC (2.31 vs 3.03 ± 0.15, respectively) and were significantly lower in LF than in NC at 14 d post-application (2.82 vs 3.61 ± 0.22, respectively), indicating that application of LF may have reduced bacterial heterogeneity on the hutch surfaces. Evenness of species distribution according to Simpson’s index was also significantly different between treatments (Treatment, *P* = 0.04). The PCoA of the Unifrac distances for beta-diversity (Figure 2.1) indicates a distinct clustering of LF treated samples after 24 hours of application compared with other treatment groups and timepoints, which was partly sustained after 7 days post-application. In this study, LF-treated hutches had higher relative abundances of ASV belonging to *Pediococcus* and *Arthrobacter* as well as reductions in ASVs belonging to *Acinetobacter*, *Corynebacterium*, *Lactococcus*, *Planococcus*, *Planimicrobium*, *Pseudomonas*, and *Streptococcus* (Figure 2).

**Figure 1:**
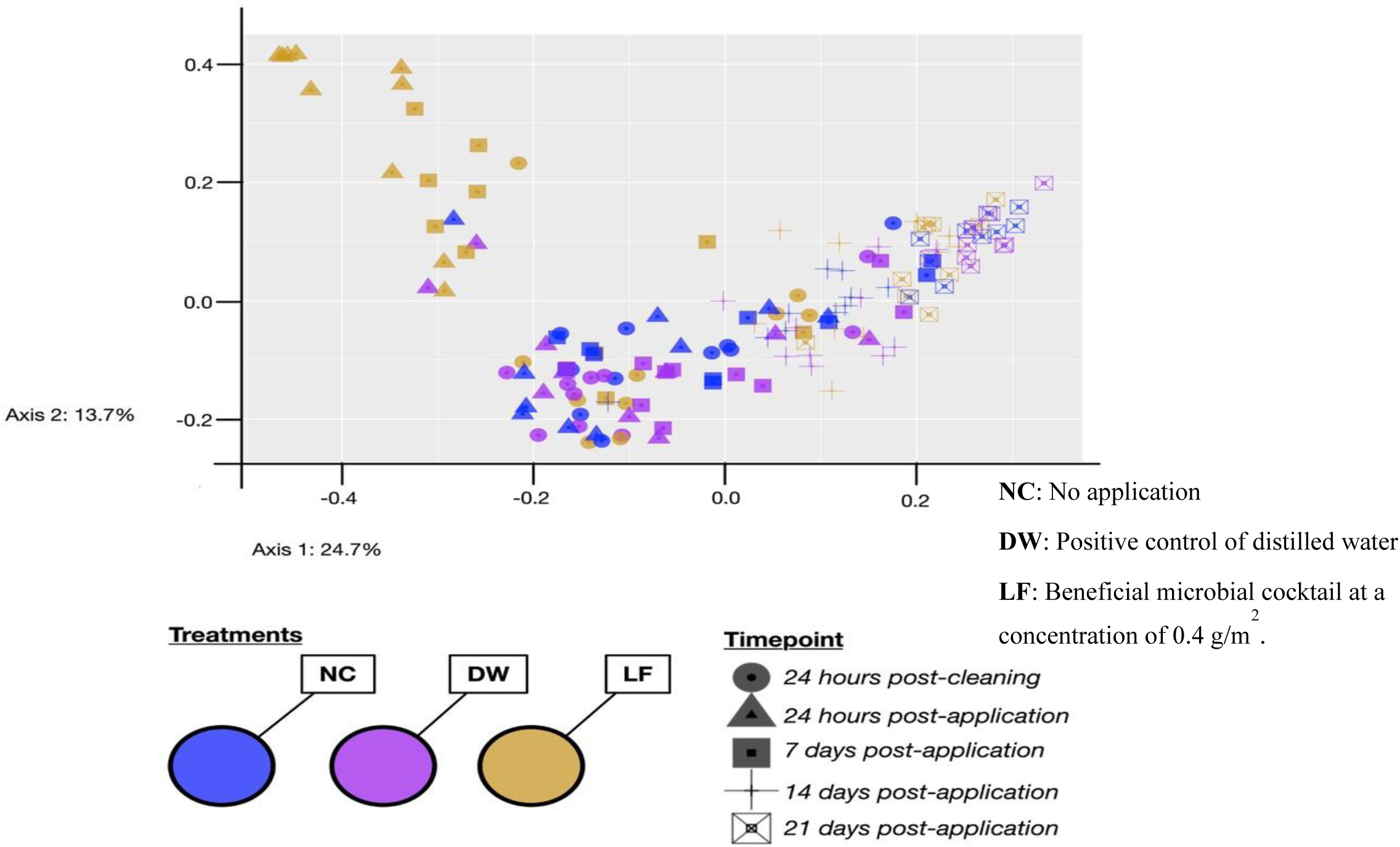
Principal Coordinates Analysis (PCoA) on the weighted UniFrac distances of samples across a 21-d treatment period from polyethylene calf hutch surfaces treated with a negative control (NC; no application), a positive control of distilled water (DW), or a beneficial microbial cocktail containing *Bacillus spp.* and *Pediococcus spp*. at a concentration of 0.4 g/m^2^ (LF).

**Figure 2.**
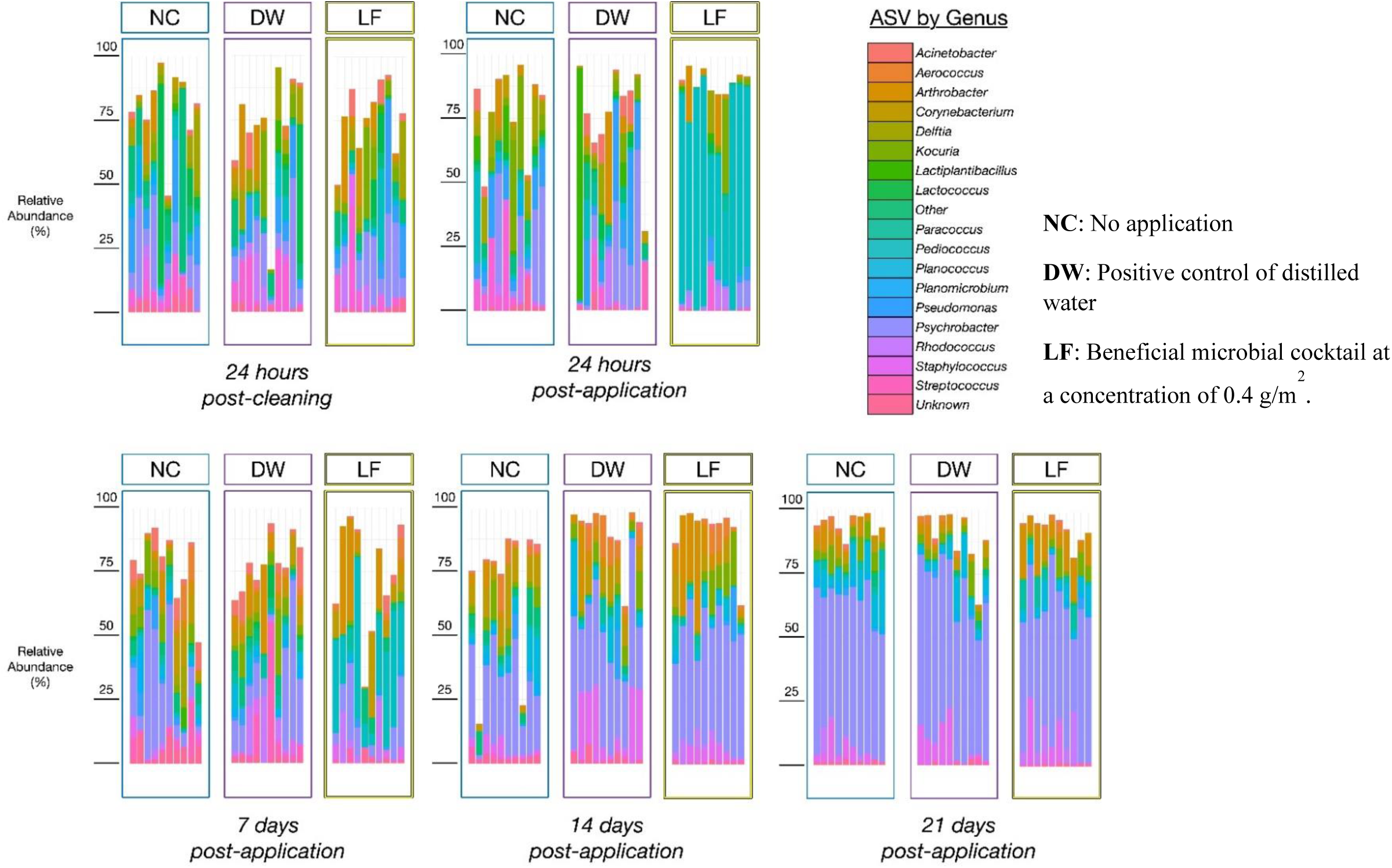
Relative abundance (%) of the 20 most abundant amplicon sequence variants (ASVs) at the genus level from polyethylene calf hutch surfaces treated with treated with a negative control (NC; no application), a positive control of distilled water (DW), or a beneficial microbial cocktail containing *Bacillus spp.* and *Pediococcus spp*. at a concentration of 0.4 g/m^2^ (LF). Each bar plot represents an individual sample (n=10/trt).

**Table 4.** Alpha-diversity indices as indicators of bacterial diversity present over a 21-d treatment period on polyethylene calf hutch surfaces treated with a negative control (NC; no application), a positive control of distilled water (DW), or a beneficial microbial cocktail containing *Bacillus spp.* and *Pediococcus spp*. at a concentration of 0.4 g/m^2^ (LF).

| Diversity Indices/Treatment<sup>1</sup> | Overall mean | Time Post-Application |  |  |  | SE | P-values |  |  |
| --- | --- | --- | --- | --- | --- | --- | --- | --- | --- |
|  |  | 24 h | 7 d | 14 d | 21 d |  | Treatment | Time | Treatment × Time |
| **Observed Species** |  |  |  |  |  | 6 | 0.005 | <0.001 | 0.14 |
| NC | 123 | 64 | 165 | 161<sup>a</sup> | 103 |  |  |  |  |
| DW | 108 | 51 | 158 | 119<sup>ab</sup> | 102 |  |  |  |  |
| LF | 91 | 56 | 119 | 95<sup>b</sup> | 94 |  |  |  |  |
| **Shannon Index** |  |  |  |  |  | 0.15 | <0.001 | <0.001 | 0.29 |
| NC | 3.29 | 3.03<sup>x</sup> | 3.85 | 3.61<sup>a</sup> | 2.65 |  |  |  |  |
| DW | 2.95 | 2.64 | 3.59 | 3.14<sup>ab</sup> | 2.45 |  |  |  |  |
| LF | 2.76 | 2.31<sup>y</sup> | 3.31 | 2.82<sup>b</sup> | 2.61 |  |  |  |  |
| **Simpson Index** |  |  |  |  |  | 0.01 | 0.04 | <0.001 | 0.09 |
| NC | 0.88 | 0.90 | 0.93 | 0.92 | 0.75 |  |  |  |  |
| DW | 0.84 | 0.85 | 0.90 | 0.84 | 0.71 |  |  |  |  |
| LF | 0.84 | 0.81 | 0.92 | 0.84 | 0.78 |  |  |  |  |

<sup>1</sup> Treatments were a negative control (NC; no application), a positive control of distilled water (DW), or a beneficial microbial cocktail (LF; Lallemand SAS, Blagnac, France) at a concentration of 0.4 g/m<sup>2</sup>.
<sup>ab</sup> Means within each column with differing superscripts were significantly different ( $P \leq 0.05$ ).
<sup>xy</sup> Means within each column with differing superscripts tended to be different ( $0.05 < P \leq 0.10$ ).

Differential abundance analysis was performed with IndicSpecies, and selected ASV’s of interest were reported (Table 5). At present, more ASVs were identified as indicator species in samples originating from NC groups compared with LF and DW groups. This included multiple ASV associated with soil and gastrointestinal origins after 14 days, and ASVs belonging to *Corynebacterium*. Furthermore, DW had some ASVs belonging to pathogens identified as indicator species including *Paeniclostridium* after 24 hours, *Staphylococcus* after 14 days, and *Escherichia-Shigella* after 21 days. Additional indicator species identified in DW included ASVs that have been identified as potentially opportunistic pathogens found in the environment and gastrointestinal tracts of humans and livestock including *Actinomyces* and *Moraxella.* In contrast, LF were consistently associated with ASVs belonging to *Bacillus* and *Pediococcus* after 24 hours post-application as well as 7- and 14-days post-application. Finally, ASV belonging to *Bacillus* was also identified as an indicator species after 21 days post-application in LF-treated hutches.

**Table 5.**
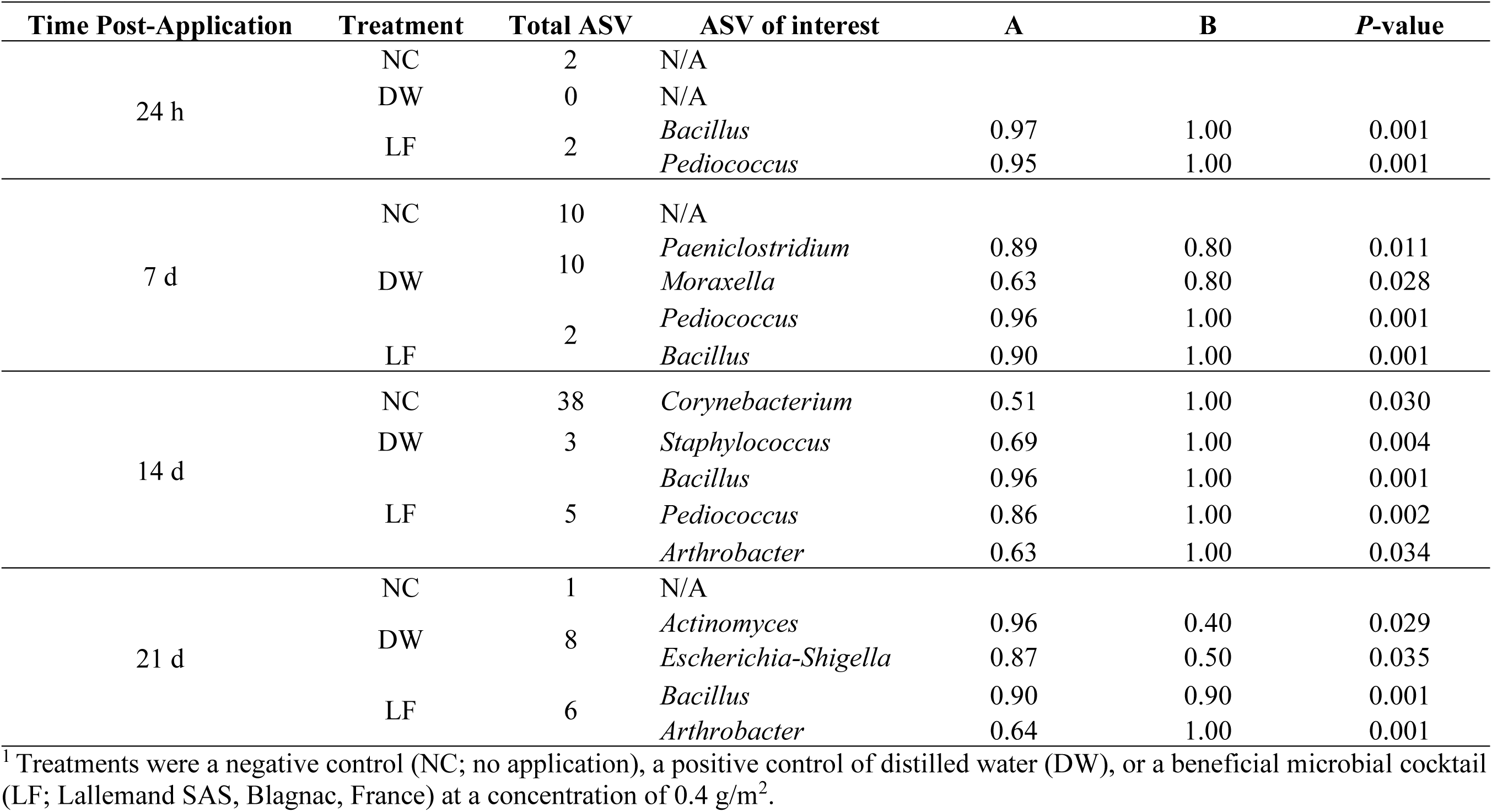
Number of amplicon sequence variants (ASVs) at the genus-level identified as indicator species, the ASVs of interest, indicator output results, and *P-*values of the relationships observed within treatment groups after 24 hours (h), 7, 14, and 21-days (d) post application on polyethylene calf hutch surfaces treated with a negative control (NC; no application), a positive control of distilled water (DW), or a beneficial microbial cocktail containing *Bacillus spp.* and *Pediococcus spp*. at a concentration of 0.4 g/m^2^ (LF).

## DISCUSSION

In pre-weaned calf housing, where sanitation is of utmost importance, the employment of beneficial biofilms may be a valuable complement to cleaning and disinfection. This is the first known use of this beneficial biofilm product in the pre-weaned calf environment, and its ability to adhere to and influence the microbial communities present on calf housing surfaces is a promising result. The microbiome of swine environments has been examined to better categorize biofilm formation and bacterial diversity in livestock environments (Guéneau et al., 2021), but a lack of detailed understanding of biofilm development mechanisms and their inhibition potential against undesirable bacteria remains. The results from this study supplement previous work by Guéneau et al. (2022a, b) by providing more data to support the use of beneficial biofilms in livestock environments as well as characterize the structure of the bacterial communities in calf housing environments using 16S amplicon sequencing. As calves often engage in non-nutritive oral behaviors with items present in their housing environment, reducing the amount of orally transmitted pathogens on surfaces would be a positive outcome.

This study was performed during the summer months to provide favorable temperatures for bacterial growth, and the laboratory conditions were set to mimic the temperature and humidity conditions of the calf environment. *E. coli* growth in the animal intestine is optimal between 36-40°C, but proliferation in the environment occurs at temperatures < 30°C (Jang et al., 2017). *Salmonella enterica* serovar Typhimurium can tolerate more harsh conditions, with growth documented in temperatures ranging from 5-45°C (Matches and Liston, 2006). *Cryptosporidium parvum* oocysts can be viable for weeks in soil and water at various temperatures, and are quite resistant to eradication (Arrowood, 2002). These three pathogens are often the causative agents of diarrhea in calves less than one month of age (Constable, 2009; Cho and Yoon, 2014), and so were chosen as targets for detection on calf housing surfaces.

Adherence and longevity of static biofilms to surfaces are dependent on several factors, such as nutrient availability, temperature, moisture, pH, surface characteristics, and existing microbial communities (Merritt et al., 2011; Bhagwat et al., 2021). Spore-forming bacteria preferentially adhere to rough hydrophobic surfaces, such as the polyethylene used to construct calf hutches (Muhammad et al., 2020; Bhagwat et al., 2021). *Bacillus* spp. typically form a pellicle at air-solid interfaces, which aids in attachment to the surface and provides a protective component to the maturing biofilm (Arnaouteli et al., 2021). Disruption of cellular mechanisms and competition for nutritional and spatial resources give LAB and other positive biofilm-forming bacteria an inhibitory advantage over other environmental bacteria (Guéneau et al., 2022b). Detection of ASVs belonging to *Pediococcus* and *Bacillus* genera supports successful product application in situ. A significant increase in the relative abundance of ASV belonging to *Pediococcus* was detected after 24 h post-application, which comprised nearly 65% relative abundance, and seems to have maintained establishment on calf hutches after 7 d post-application at around 25% relative abundance. Although 16S sequencing is unable to discriminate between the LAB and *Bacillus* present in the product from those naturally present in the environment, taxa belonging to *Pediococcus* and *Bacillus* were identified as indicator species in LF-treated hutches. Further analysis needs to be conducted to determine if, and when, re-application of the product may be necessary.

The Indicator Value index was defined as a means to measure associations between different ‘sites’ and species present for ecological studies (Dufrene and Legendre, 1992). In the context of using Indicator Values for microbial community datasets, such analysis can identify bacterial populations that are not only distinct to different environments/ experimental groups but can also indicate the extent of their prevalence within groups. In this study, taxa belonging to *Pediococcus* and *Bacillus* genera were not only indicated as being distinct in LF-treated hutches (A-values > 0.85 for 24 h, 7, and 14 d post-application), but their prevalence within each group was very confident (B-values = 1.00 for 24 h, 7, and 14 d post-application, indicating that these taxa were observed in all samples from LF group). Finally, taxa belonging to *Bacillus* after 21 d post-application were identified as being distinct to LF-treated hutches (A = 0.90) and prevalent across most samples (B = 0.90) which could suggest prolonged establishment of *Bacillus* post-application. Finally, Supplementary Figure 2 depicts a comparison of bacterial colonies enumerated on TSA from samples taken from both NC and LF hutches after 24 h post-product application indicating a visual difference in bacterial heterogeneity between the two treatments. Although this result is observational in nature, the wrinkled pellicle colony formation aligns with the physical characteristics of most *Bacillus* spp. in culture and is clearly dominant in the sample from the LF hutch after 24 h post product application compared to the diversity in colony morphologies present in the sample from the NC hutch.

The primers chosen for the detection of *Escherichia* spp.*, Salmonella spp.,* and *C. parvum* from in situ samples were based on target genes present in most or all the selected pathogens. The *uidA* gene that encodes for β-glucuronidase is present in 94-96% of all *E. coli* strains (Perin et al., 2010; Miotto et al., 2019). The invA gene is successfully detected in all *Salmonella* serovars (Nurjanah et al., 2018), and the COWP gene responsible for *Cryptosporidium* oocyst wall integrity is also a common qPCR target gene (Guy et al., 2003). It is important to note that this method did not utilize PMA-mediated DNA extraction, and therefore is unable to distinguish between DNA from live or dead cells (Gensberger et al., 2014). Because DNA concentrations for all three pathogen targets were below the detection threshold of 3 log copies, these results are inconclusive regarding definitive interactions between, and effects of, beneficial biofilm application on competing microbial communities.

Alpha diversity at the within-habitat level summarizes the structure of ecological communities with respect to their overall taxonomic richness and distribution of abundances (Thukral, 2017; Willis, 2019). The Shannon-Weaver and Simpson’s indices have historically been the most robust measure of microbial diversity (Kim et al., 2017). The decrease in species richness in LF-treated hutches aligns with the higher relative abundance of *Pediococcus* and other LAB species detected, potentially indicating their domination of the surface ahead of other competing communities and reducing the heterogeneity of bacterial species present in the sample. Similar results were observed from samples recovered from a poultry broiler barn treated with a positive biofilm compared to an untreated barn (Guéneau et al., 2022a). The diversity results in our study are also further supported through the Indicspecies outputs. For instance, the total taxa identified as indicator species (A values ≥ 0.50) in NC hutches were 10 and 38 at 7 and 14 d post-application, respectively, compared to 2 and 5 ASVs in LF treated hutches. None of the taxa in LF-treated hutches were pathogenic or opportunistic pathogens.

Use of beneficial bacteria have been implicated as potential OneHealth approaches to disease management on farms (Guéneau et al. 2022a, 2022b). Exposure to certain bacteria can impact the future performance of calves and the health of agricultural workers. In this study, reductions were observed in taxa belonging to zoonotic bacterial species in LF-treated hutches including *Streptococcus* spp., *Corynebacterium* spp., and *Pseudomonas* spp. Reductions in taxa belonging to *Streptococcus* in LF-treated hutches is a promising finding, as this genus is implicated as a mastitis pathogen and is commonly found in manure and animal bedding (Alanis et al., 2021). Duniere et al. (2024) also observed decreases in *Streptococcus* spp. when a beneficial bacterial cocktail was applied to recycled manure solids used as bedding for lactating cows.

LF-treated hutches also demonstrated reductions in the taxa belonging to *Corynebacterium,* which are commonly identified in agricultural environments. Furthermore, this taxa was identified as an indicator species in NC hutches after 21 d post application. *Corynebacterium* spp. have also been identified as mastitis-causing pathogens and are frequently isolated from bovine mammary glands (Hogan et al., 1988; Watts et al., 2001; Dalen et al., 2018). While *Corynebacterium* spp. have not been known to cause severe disease in calves, some strains can cause diphtheria-like infections in humans working in close contact with animals (Bernard, 2012). However, certain *Corynebacterium* spp. have recently been identified as potential human probiotics with similar mechanisms to lactic acid bacteria for antioxidant potential and promoting gut health (Shamsuzzaman et al., 2023), so there may be other enzymatic and metabolic interactions with LAB on the surface that could be explored in future work.

*Pseudomonas* spp. are known biofilm formers and opportunistic pathogens frequently associated with foodborne illness outbreaks, gastrointestinal illness, milk spoilage, and mastitis in cattle (Van Tassel et al., 2012). *Pseudomonas aeruginosa,* the most clinically important strain, is ubiquitous in water, soil, and the urogenital microflora of healthy cows (Giannattasio-Ferraz et al., 2022). Given their propensity toward wet environments, it is not surprising to observe a greater relative abundance of *Pseudomonas* spp. in those hutches treated with a positive control of distilled water. Humid areas and soiled, wet bedding are prime environments for growth and biofilm formation of *Pseudomonas* spp. (Schauer et al., 2021; Badawy et al., 2023), and the relative humidity observed in the calf environment during this study would support its proliferation. Antimicrobial resistance and biofilm-forming capacity of *Pseudomonas* spp. has attracted the focus of the World Health Organization, designating them as “priority pathogens” for new antimicrobial targets (WHO, 2017). Certain LAB, particularly *Pediococcus* spp., have demonstrated strong antagonistic effects against *Pseudomonas* spp. Metabolites from a strain of *Pediococcus pentosaceus* isolated from cow’s milk disrupted the quorum sensing mechanisms required for biofilm formation by *Pseudomonas aeruginosa* isolated from rancid butter (Aman et al., 2021). These previous findings support the reductions of taxa belonging to *Pseudomonas* genus in LF-treated hutches. Many *Bacillus* species produce volatile and antimicrobial compounds, which interfere with cell-cell signaling and allow them to inhibit or outcompete biofilm production by neighboring bacteria (Moore et al., 2013; Caulier et al., 2019; Hou et al., 2021). However, other findings with *Bacillus* spp. support an alternative theory. More recent work has also explored what Lyng and Kovács (2023) describe as a ‘frenemy’ relationship between certain *Bacillus* spp. and *Pseudomonas* spp., suggesting that their coexistence in the environment results in amensalism, therefore disrupting or disabling each species’ cellular and metabolic responses for protection or spatial competition. Despite the use of *Bacillus spp.* in the beneficial biofilm-forming cocktail, our results indicate there may have been exclusionary behavior toward *Pseudomonas* spp. perhaps due to the inclusion of LAB strains *Pediococcus* spp. in the formula.

Increases in taxa belonging to the genus *Arthrobacter* in LF treated hutches may insinuate impacts to resources rather than animal health. This genus has been identified on polyethylene surfaces and has been implicated in degradation of this type of plastic (Han et al., 2020). This emphasizes the necessity of good cleaning and sanitation practices to remove soil from surfaces, further solidifying that the use of competitive-exclusion bacteria are a complement to, not a substitution, for cleaning and disinfection procedures (Luyckx et al., 2016). Especially considering this ASV in the present study was identified as an indicator specie for LF-treated hutches after 14 and 21 d post application (A= 0.60 and 0.64, B = 0.80 and 1.00 respectively), additional research efforts could focus on understanding the synergies and potential antagonistic interactions between positive biofilm-forming species and *Arthrobacter* spp. to reduce risk for degradation of agricultural plastics.

## CONCLUSIONS

A beneficial bacterial cocktail of *Bacillus* spp. and *Pediococcus* spp. was able to successfully colonize on polyethylene, and its application to individual polyethylene calf housing influenced microbial diversity and reduced the presence of undesirable bacteria on high-contact interior surfaces. This is a novel approach to biosecurity in the pre-weaned calf environment and may prove a valuable complement to cleaning and disinfection procedures to diminish pathogen load on surfaces. Further studies with varied microbial load intensities and different housing and management strategies are necessary to elucidate antagonistic and amensalistic interactions between bacterial communities present in calf housing, potential health outcomes, and broader applicability in calf-rearing operations.

## ACKNOWLEDGEMENTS

This study was funded by Lallemand SAS (Blagnac, France). Authors Richard Scuderi, Andrew Skidmore, and Lysiane Duniere were employed by the Animal Nutrition Business Unit of Lallemand Inc. when the study was performed. However, their affiliation did not impede their ability to follow journal guidelines or to remain impartial during the preparation of the manuscript. Gratitude is extended to Dr. Pascal Drouin (independent researcher) for technical assistance during the study, and to Dr. Virgile Guéneau for editorial assistance.

## DATA AVAILABILITY

The 16S sequence datasets generated during this current study are available on NCBI under accession number PRJNA1188585.

---

**Supplementary Figure 1.**
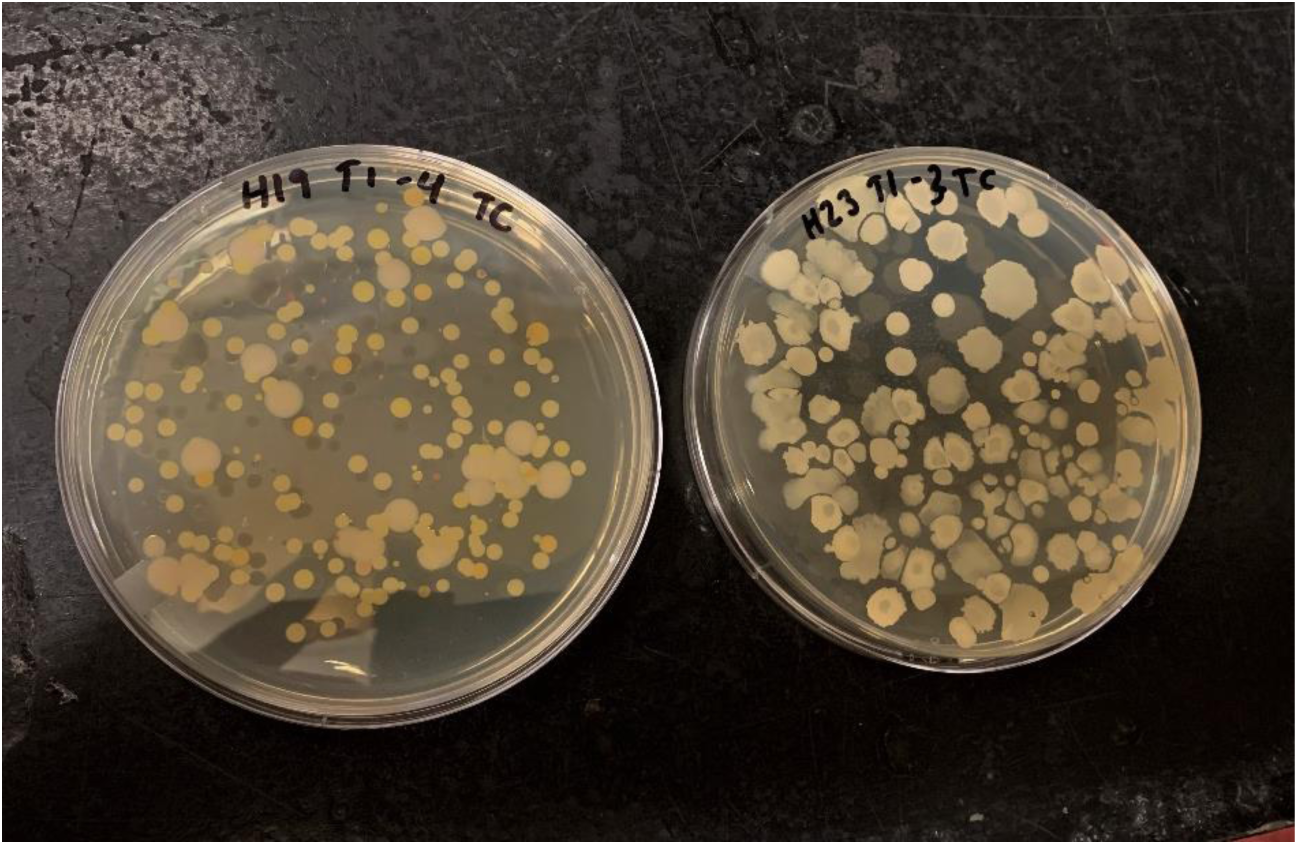
Total cell counts on tryptic soy agar enumerated from samples obtained 24 h post-product application from polyethylene calf hutches treated with either a negative control (NC: no application) or a beneficial microbial cocktail (LF; Lallemand SAS, Blagnac, France) at a concentration of 0.4 g/m^2^.

**Supplementary Table 1.**
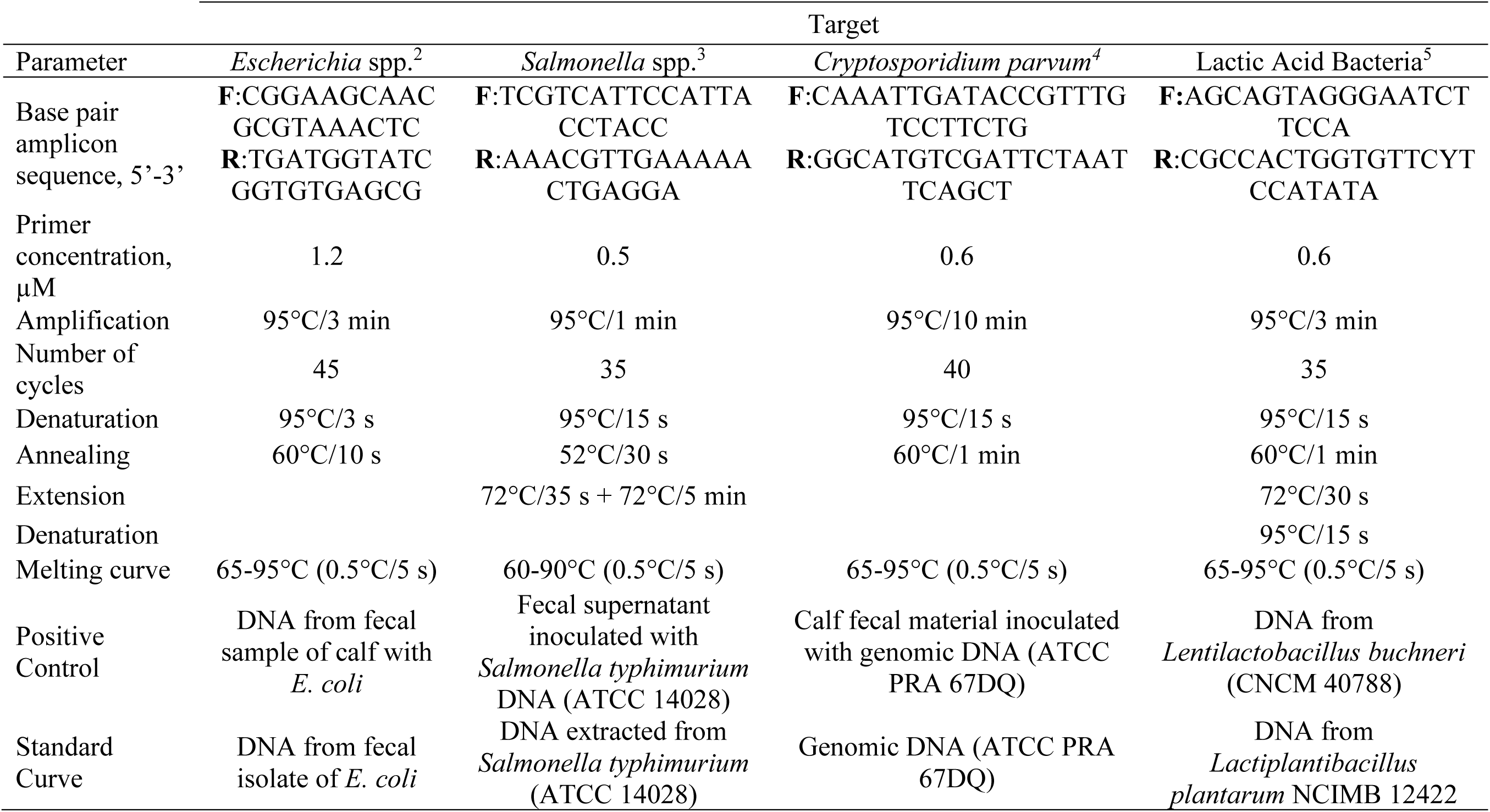
Parameters of RT-qPCR analysis performed to detect microbial DNA in situ over a 21-d treatment period^1^ on polyethylene calf hutch surfaces treated with treated with a negative control (NC; no application), a positive control of distilled water (DW), or a beneficial microbial cocktail containing *Bacillus spp.* and *Pediococcus spp*. at a concentration of 0.4 g/m^2^ (LF).

